# Unifying evolutionary dynamics: a set theory exploration of symmetry and interaction

**DOI:** 10.1101/2023.09.27.559729

**Authors:** Francisco Berkemeier, Karen M. Page

**Affiliations:** Department of Genetics, University of Cambridge, CB2 3EH, Cambridge, United Kingdom; Department of Pathology, University of Cambridge, CB2 1QP, Cambridge, United Kingdom; Department of Mathematics, University College London, WC1E 6BT, London, United Kingdom; Institute for the Physics of Living Systems, University College London, WC1E 6BT, London, United Kingdom

**Keywords:** Evolutionary Dynamics, Interaction Kernels, Symmetry, Axiom of Choice

## Abstract

Within the expansive landscape of evolutionary dynamics, symmetry features embedded in well-established models significantly influence the interpretation of individual interaction patterns. Such symmetries are determined through interaction kernel functions, which serve as mathematical models for characterizing the complexity of interactions between individuals, each with distinct phenotypes. By incorporating analytical tools from logic and set theory, we aim to provide a deeper understanding of these functions, relevant to mechanisms of evolution. We prove that the kernels introduced in Champagnat et al.’s unifying framework exist provided birth and death rates are symmetric with respect to non-focal traits. The kernels may nevertheless be highly challenging to construct, thereby indicating a complex underlying mathematical infrastructure within unified evolutionary dynamics. We show how interaction kernels for asymmetric frameworks arising in evolutionary graph theory can be derived by incorporating individuals’ graph labels into their phenotypes. These insights invite new avenues for research, providing a fresh understanding of the interactions between individuals in broader biological contexts.

## Introduction

In the mid-19th century, Charles Darwin put forth the transformative theory of natural selection, suggesting that species undergo changes over time (Darwin et al., 1958; Darwin, 1987). This theory posits that each species possesses unique, inheritable genetic variations that have slowly evolved from a shared ancestor over a vast period of time, ultimately constructing an intricate tree of life. This tree manifests through a continual and varied sequence of branching events, including reproduction, death, and mutation, giving birth to new species in a phenomenon referred to as speciation. These fundamental processes permeate all of biology. Evolution shaped by natural selection promotes advancement and enhancement. More specifically, the traits that contribute to increased fitness within a species’ phenotype are likely to become more preva-lent. Darwinian fitness can be loosely understood as the capacity of an individual to live long enough to procreate successfully numerous times in order to sustain the population or species.

In early debates, biometricians W. F. R. Weldon and Karl Pearson presented arguments in favour of Darwin’s theory, asserting that even minor variations could wield considerable influence on evolutionary progress (Darwin, 1859; Martins, 2007). In contrast, Francis Galton and William Bateson proposed that evolution fundamentally occurs in leaps and bounds, facilitated by the sudden appearance of markedly different individuals capable of passing on their phenotypes to future generations (Norton, 1973; Olby, 1989). As the scientific understanding evolved, the concepts of Mendelian genetics and Darwinian evolution were harmoniously integrated by Haldane, Fisher, and Wright (Haldane, 1956; Fisher et al., 1958; Haldane and Jayakar, 1963; Wright, 1969). This synthesis expanded the mathematical framework for modelling and interpreting these systems. Such mathematical articulation of evolutionary dynamics laid the foundation for groundbreaking theories, including Hamilton’s kin selection (Hamilton, 1970), Kimura’s theory of neutral evolution (Kimura and Harper, 1970), and Maynard Smith’s evolutionary game theory (Smith, 1982). Overall, the cornerstone of mathematical models for adaptation and co-evolution of biological populations within Darwinian dynamics, focusing on selection and mutation, lies in a comprehensive understanding of population genetics combined with evolutionary game theory.

Processes that enhance fitness are typically instrumental in shaping evolutionary dynamics by posing an optimisation problem amenable to modelling. Over time, these dynamics often reach a predictable equilibrium after resolving the problem at hand. However, stochastic effects arise in these systems due to genetic mutations and environmental changes. To analyse frequencydependent selection across the phenotypic space, the integration of evolutionary game theory becomes an essential part of the mathematical and computational approach to biology.

The triad of evolution – reproduction, mutation, and selection – has been the subject of extensive research within the domain of evolutionary dynamics. Notably, a number of studies have attempted to integrate various evolutionary dynamics modelling methodologies in multiple ways. A unified framework for modelling evolutionary processes was created in the deterministic scenario, taking into account a continuous relative fitness function of strategically interacting individuals. Here, the authors examined broad categories of evolutionary dynamics and compared them to Nash equilibria (Kreps, 1989; Joosten, 1996). Leveraging the equivalence between the replicator-mutator equation, which delineates the dynamics of population distribution (Hadeler, 1981), and the Price equation, describing its moments (Price et al., 1970; Price, 1972), numerous mathematical models of evolutionary dynamics were demonstrated to fall under a cohesive, unified framework (Page and Nowak, 2002). These equations have been revealed to spawn a multitude of modelling contexts, such as adaptive dynamics (Nowak and Sigmund, 1990; Metz et al., 1995; Dieckmann and Law, 1996), evolutionary game dynamics (Hofbauer et al., 1998), the Lotka-Volterra ecological equation (Volterra, 1926), and the quasi-species equation pertinent to molecular evolution (Eigen et al., 1989). Evolutionary dynamics were also studied under different timescales and rescaling limits by N. Champagnat and colleagues (Champagnat et al., 2006). It was discovered that the architectural framework of macroscopic models within evolutionary dynamics is considerably shaped by the temporal aspects of individual events. In this context, the authors presented a detailed model that encapsulates the stochastic dynamics of birth, mutation, and death in continuous time, influenced by each individual’s trait values and their interactions with others. This culminated in a general algorithm for an efficient numerical simulation of the individual-level model. Using the stochastic point process, an array of macroscopic models of adaptive evolution was generated, each differing based on the assumed renormalisation (such as population size, mutation rate, and mutation step), thereby influencing their deterministic or stochastic nature under precise time rescaling.

Early in the discussion of this unifying description of evolutionary dynamics, microscopic population point processes were introduced as dependent on the interaction between different individuals, characterised by their phenotypes. Such a dependence was written in terms of interaction kernels, which provide the basis of the process construction. Our study aims to identify the constraints of such a modelling framework. Specifically, we strive to comprehend how symmetry with respect to (w.r.t.) non-focal phenotypes, reliant on these interaction kernels, is upheld within this framework. Our investigation leads us to a deep theoretical discourse, resting on the principles of the Axiom of Choice and Zorn’s Lemma. Although the biological interpretation of the Axiom of Choice offers a fascinating philosophical topic on its own, it falls outside the purview of this work.

We revisit the point process modelling notions in Champagnat et al. (2006), along with the primary instruments from logic and set theory employed in validating the central theorems in this work. Ultimately, we highlight some implications of these theorems with examples concerning evolutionary games defined on graphs.

## Theoretical framework

Here we present the main theoretical background, including the relevant concepts presented in Champagnat et al. (2006), as well as the set theory elements essential to the main discussion.

### Evolutionary dynamics

The explanation of the population point process in Champagnat et al. (2006) makes use of several mathematical techniques and ideas. The comprehensive exposition below closely follows the discussion presented in that paper.

Each individual’s phenotype is represented as a real-valued vector of trait values. The trait space *X* satisfies *X* ⊆ ℝ^*ℓ*^, where *ℓ* is the number of traits. A counting metric that counts the number of individuals displaying various phenotypes is used to characterise the population as a whole, evolving according to a Markov process (Gagniuc, 2017). The Markov property presupposes that the population’s dynamics at time *t* depend exclusively on the population’s current state.

Here, we take into account a population in which individuals can reproduce and die at rates that are determined by their unique traits as well as by their interactions with others who have similar or different traits. These events occur continuously and at random. Reproduction is almost faithful, but there is a chance that a mutation will result in an offspring with a phenotype that is different from that of its ancestor. For the purpose of our discussion, however, we do not account for mutations.

Birth and death rates of focal individuals with phenotype *x*∈ℝ^*ℓ*^ are defined as functions of *x* and their interactions with other individuals. These interactions are measured by an *interaction kernel*, which is a function of how phenotypically different they are. For a population of size *n*, this dependence is explicitly given by

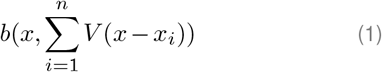

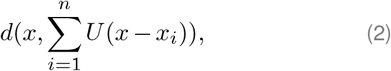

where *b* and *d* are the birth and death rates, respectively, and *V* and *U* are the interaction kernels affecting reproduction and mortality. We may now question the generality of this approach and which ways of defining such rates are not being considered by writing the dependence as above.

Symmetry w.r.t. non-focal phenotypes is a first property that is worth discussing. Indeed, both birth and death rates with trait dependence given by Eq. (1)-Eq. (2) are symmetric w.r.t. non-focal traits, which has intricate implications derived from set theory that we explore in the following sections.

### Elements of set theory

We are interested in determining what are the properties of a real function *h* such that there exist functions *f* and *g* satisfying

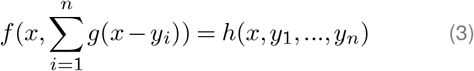

for all *x, y*_1_,.., *y*_*n*_∈ℝ^*ℓ*^ and *n*≥2. We claim that Eq. (3) holds, for some functions *f* and *g*, if and only if *h* is symmetric w.r.t. *y*_1_, …, *y*_*n*_, that is, its value is the same independently of the order of its arguments. For example, a function *f* (*x*_1_, *x*_2_) taking two arguments is a symmetric function if and only if *f* (*x*_1_, *x*_2_) = *f* (*x*_2_, *x*_1_) for all *x*_1_ and *x*_2_ such that (*x*_1_, *x*_2_) and (*x*_2_, *x*_1_) are in the domain of *f*. More relevant to our case, we can see that a function of the form

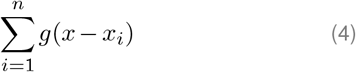

is also symmetric w.r.t. *x*_1_, …, *x*_*n*_. To begin the thought process behind the main theorems in this work, we present some examples with *n* = 2.

#### Example 1

(A functional equation). Suppose

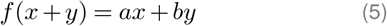

holds for some function *f* : ℝ→ ℝ and *a, b* ∈ ℝ. Given the function 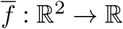 defined as 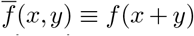 we see that it is symmetric w.r.t. *x* and *y*, since *x* + *y* = *y* + *x*. Then, it follows that

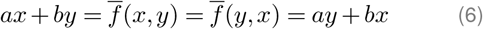

and thus *a* = *b*. Hence, if *a* ≠ *b*, Eq. (5) does not hold for any function *f*. In some sense, symmetry imposed by 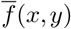 is extended to *ax* + *by*. A more straightforward way to see this is by noting that *f* (0 + 0) = *f* (1−1) implies *a* = *b*.

#### Example 2

(Symmetry necessity). Let *h*(*x, y, z*) = *ax* + *by* + *cz*, where *a, b, c* ∈ℝ. Then Eq. (3) becomes

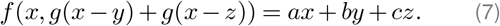

However, from Example 1, 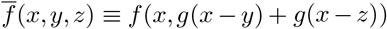 is symmetric w.r.t. *y* and *z* and so

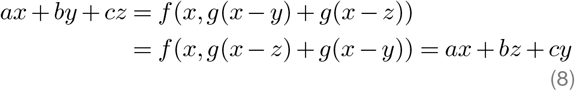

thus *f* and *g* exist if and only if *b* = *c*. In particular, take *g*(*w*) = *w* and *f* (*u, v*) = *au* + *b*(2*u*−*v*). For *n >* 2, the reasoning is similar. This example aims to suggest that some kind of symmetry in *h* in Eq. (3) is necessary. In fact, since *f* is symmetric w.r.t. *y*_1_, …, *y*_*n*_, *h* is neces-sarily symmetric w.r.t. the same variables, thus we now focus on the reverse implication.

#### Example 3

(Symmetry sufficiency). We now illustrate that symmetry of *h* might not only be necessary, but also sufficient to guarantee the existence of *f* and *g* such that Eq. (3) holds. Since isolated dependence on *x* is granted from *f*, we drop it from *h* and assume the non-linear case *h*(*y, z*) = *y*^2^ + *z*^2^, which is symmetric w.r.t. *y* and *z*. Thus

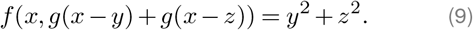

Here, the answer is not trivial and further machinery is needed. With a suitable change of variables, we can rewrite Eq. (9) as

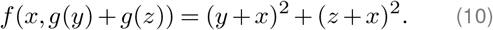

Since we cannot explicitly impose dependence of *g* on variable *x*, the only way to satisfy Eq. (10) is to find a function *g* such that the sum *g*(*y*) + *g*(*z*) uniquely determines the unordered pair (*y, z*). This would then guarantee explicit dependence of *f* on (*y, z*), simplifying the problem. Note that the unordered property is a consequence of symmetry. Ideally, if *y, z* ∈ℝ, we would like *g* to be a function between ℝ and any set with the cardinality of the continuum (cardinality of ℝ, usually denoted by 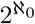), such that any two sums in that set give distinct values. Such a set and bijection are not trivial to construct and their existence relies on Zorn’s Lemma, discussed next.

### On the existence of a basis and the Axiom of Choice

By interpreting the real numbers as a vector space over some set, constructing a basis for ℝ and defining *g* in Eq. (10) as the bijection between ℝ and its basis leads to the unique description of the unordered pair (*y, z*), since *g*(*y*) and *g*(*z*) are linearly independent over the set we build our basis on. In general, this would then allow us to take the right-hand side of Eq. (3) as any symmetric function w.r.t. non-focal traits.

One way to do this is to consider ℝ as a vector space over the rationals, where a basis is a collection ℬ of real numbers such that every real number is a rational linear combination of numbers *β*_*i*_ ∈ ℬ in precisely one way, that is, every real number *α* has a unique representation of the form

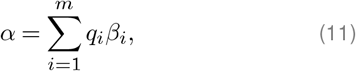

where *q*_*i*_ is rational and *m* depends on *α*. This set is also known as the *Hamel basis* for ℝ(Hamel, 1905). Note that any Hamel basis contains, at most, one rational number (Figure 1a). Considered as a vector space over ℚ, ℝ has uncountable dimension, so it cannot be spanned by countably many vectors. Hamel bases have been studied in the context of additive functions (Jones,1942; Boros and Daróczy, 2006), including the discussion of the general solution of Cauchy’s functional equation (Aczél and Erdő s, 1965). The existence of such a basis depends on the *Axiom of Choice* or, more directly, *Zorn’s Lemma* (Zermelo, 1904; Kuratowski, 1922; Zorn, 1935). In the following, we are then considering an axiomatic system under Zermelo-Fraenkel set theory with the Axiom of Choice (ZFC) (Zermelo, 1908; Fraenkel et al., 1973).

**Figure 1.**
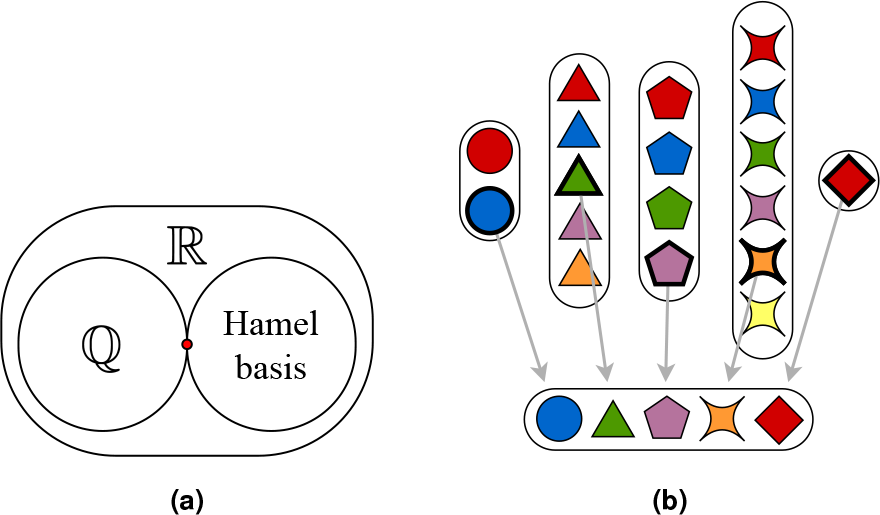
Hamel bases and Axiom of Choice. **(a)**Hamel bases contain, at most, one rational number (in red). **(b)** The choice function takes exactly one element from each set ^*α*^ (Yorgey, 2014).

The Axiom of Choice states that for any family 𝒜 of nonempty sets, there exists a set that contains exactly one element in common with each of the nonempty sets. In other words, for any family 𝒜 of nonempty sets, there exists a *choice function*

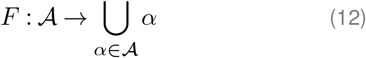

such that for every *α* ∈ 𝒜, *F*(*α*)∈ *α* (Figure 1b). These statements are equivalent to a third “version” of the Axiom of Choice:

### Zorn’s Lemma

(Axiom of Choice). *If X is a partially ordered set such that every chain in X has an upper bound, then X has a maximal element*.

It is a known result that Zorn’s Lemma, which relies on the assumption of the Axiom of Choice, yields that every vector space has a basis (Halpern, 1966; Bleicher, 1967). In the Supplementary Information, we sketch the proof of that result for ℝ.

Going back to Example 3, with *g* as the bijection between ℝ and its Hamel basis ℬ, we have that (*g*(*y*), *g*(*z*)) ≡ (*y*_*ℬ*_, *z*_*ℬ*_) ∈ ℬ ×ℬ. Hence, since B is a basis of R, *y*_*ℬ*_ and *z*_*ℬ*_ are linearly independent over the rationals and the sum *g*(*y*) + *g*(*z*) = *y*_*ℬ*_ + *z*_*ℬ*_ uniquely defines a real number. Thus it is possible to rewrite the dependence *f* in Eq. (10) as

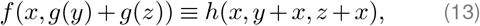

which is symmetric w.r.t. *y* and *z*. A numerical formulation of *f* and *g* is discussed later in Example 4, where we explicitly solve Example 3. Next, we present the main implications of this argument.

## Results

Recall the general phenotype-dependent expressions of birth and death rates in Champagnat et al. (2006),

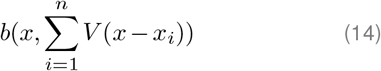

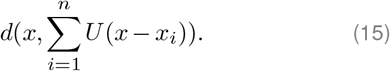

Motivated by the discussion in the previous section, we present the main theorems on symmetry derived from a phenotype-dependence like Eq. (14)-Eq. (15).

### Main theorems

Under the Axiom of Choice and via a set theory argument, one can show that the birth and death rates can be taken to be any symmetric functions.

#### Theorem 1 (General symmetry)

*Assume the Axiom of Choice. For all x, x*_1_,.., *x*_*n*_ ∈ ℝ^*ℓ*^ *and n* ≥ 2, *there exist functions f and g such that*

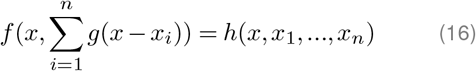

*holds if and only if h is symmetric w*.*r*.*t. x*_1_, …, *x*_*n*_.

*Proof*. (⇒) First, assume that, for some functions *f, g* and *h*, Eq. (16) holds for all *x, x*_1_, …, *x*_*n*_ ∈ ℝ^*ℓ*^. Since *f* is symmetric w.r.t. *x*_1_, …, *x*_*n*_, *h* is symmetric w.r.t. the same variables. (⇐) Now, take *ℓ* = 1 for simplicity and let *h* be any symmetric function. Considering a suitable change of variables, rewrite Eq. (16) as

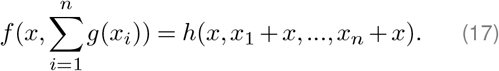

Let *g* be the bijection between ℝ and its Hamel basis, ℬ, which exists according to Zorn’s Lemma and has the cardinality of the continuum. Then,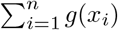 uniquely determines the unordered *n*-tuple (*x*_1_, …, *x*_*n*_), since {*g*(*x*)}_*x*_ are linearly independent over ℚ. Hence, with 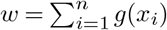, define

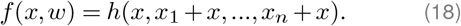

Reverting the change of variables, we get that Eq. (16) holds for all *x, x*_1_, …, *x*_*n*_ ∈ℝ. A similar argument yields the same result for any dimension *ℓ*. ◼

According to Theorem 1 and by interpreting *f* and *g* as the event rate and respective interaction kernel, Champagnat’s approach is only considering birth and death rates which are symmetric w.r.t. the trait values of the non-focal individuals, leaving a plethora of cases to be explored. We now ask which of them could be of relevant interest to evolutionary dynamics. Furthermore, the highly non-constructive nature of the proof of symmetry sufficiency (⇐) may be rather uninteresting to the applied mathematician. Therefore, in addition to the previous limitation, Champagnat’s framework is limited to a very particular type of symmetry.

### Constructible symmetry

A way to avoid the non-constructive scenario and the Axiom of Choice is to relax slightly our phenotype requirements. If we consider the traits to be vectors of rational numbers, a more direct approach may be taken. Indeed, from a computational perspective, all we really manage in simulations are approximations to real numbers, which can be seen as rational numbers (to machine error, in non-symbolic mathematics (Zazkis, 2005; Obersteiner and Hofreiter, 2017; Kristiansen, 2017; Georgiev et al., 2021)). Therefore, on rational domains, we have a weaker but constructible version of Theorem 1.

#### Theorem 1 (Constructible symmetry)

*For all x, x*_1_,.., *x*_*n*_ ∈ ℚ^*ℓ*^ *and n*≥2, *there exist constructible functions f and g such that*

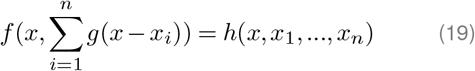

*holds if and only if h is symmetric w*.*r*.*t. x*_1_, …, *x*_*n*_.

*Proof*. (⇒) Similar to the proof of Theorem 1. (⇐) Take *ℓ* = 1 for simplicity. Considering the change of variables Eq. (17), we are now interested in finding a constructible function *g* such that 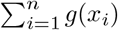 is uniquely defined for each *n*-tuple (*x*_1_, …, *x*_*n*_), i.e, such that the sum is injective w.r.t. the same tuple. Take *γ* : ℚ→ ℕ to be the bijection between ℚ and ℕ. We avoid determining an explicit analytical formula for *γ*, arguing instead that this is directly programmable with a Cantor pairing function (Szudzik, 2006; Lisi, 2007). If 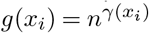, the sum

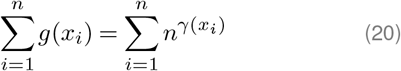

is unique for each unordered tuple of integers (*γ*(*x*_1_), …, *γ*(*x*_*n*_)) and thus for (*x*_1_, …, *x*_*n*_). To see this, let *k*_*i*_ ≡ *γ*(*x*_*i*_) ∈ ℕ. We are then looking for an injective function *σ* : ℕ → ℕ^*n*^ such that

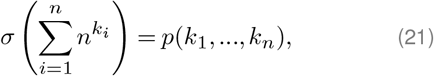

where *p* : ℕ^*n*^→ℕ^*n*^ is any permutation of (*k*_1_, …, *k*_*n*_). Then, due to the uniqueness of base-*n* representations, we have that such *σ* exists. In particular, consider the number 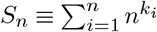 written in base-*n*, i.e,

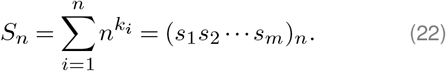

Assume first that {*k*_*i*_} arenot all equal to a specific value. Then, we have that 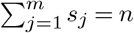 and the number *s*_*j*_ gives precisely the number of terms with exponent *j* = *k*_*i*_ that appear in 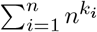. If, however, the sum is itself a power of *n* (*k*_*i*_ = *k*, ∀*i*, for some *k*), then we take *k*_*i*_ = log_*n*_(*S*_*n*_*/n*), ∀*i*. Hence, we have a unique description of the *n*-tuple via the sum. Writing *f* in terms of *σ* and applying *γ*^*−*1^ to *p*(*k*_1_, …, *k*_*n*_) implies an explicit dependence on the tuple (*x*_1_,.., *x*_*n*_) ∈ℚ^*n*^. A similar argument yields the same result for any dimension *ℓ*. ◼

Following Theorem 2, a natural extension would be to consider any countable subset of ℝ, beyond ℚ. For example, the subset of real algebraic numbers, which is countable and contains ℚ, could be of particular interest, since these represent roots of non-zero polynomials with rational coefficients, which may appear when modelling certain traits. The previous argument would then rely entirely on the constructible nature of the bijection between any countable set and ℕ. For computational purposes, however, rational traits seem to be enough.

### Further examples

#### Example 4

(Solving Example 3). Consider the one-trait scenario (*ℓ* = 1) in a population with three individuals. Let *h* : ℚ→ ℝ be defined as *h*(*x, y, z*) = *y*^2^ + *z*^2^. We now provide explicit functions *f* and *g* such that

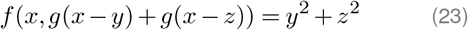

for all *x, y, z* ∈ ℚ. Again, a suitable change of variables allows us to rewrite the previous identity as

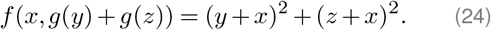

Let *γ* : ℚ → ℕ be the bijection between ℚ and ℕ, *g*(*x*) = 2^*γ*(*x*)^ and fix (*k*_1_, *k*_2_) ≡ (*γ*(*y*), *γ*(*z*)). We aim to find an injective function *σ* : ℕ → ℕ^2^ such that

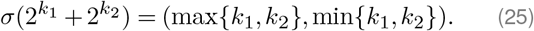

Notice first that

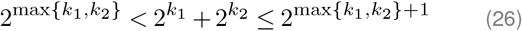

implies

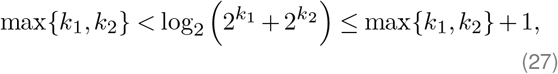

and so 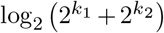falls between two successive integers. Hence, with 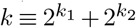 and *k*_1_ ≠ *k*_2_, we have

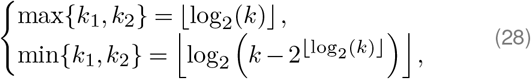

where ⌊·⌋ is the floor function, that is, ⌊*x*⌋ gives the greatest integer less than or equal to *x*. If *k*_1_ = *k*_2_, we have that 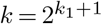 and thus log_2_ (*k/*2) = *k*_1_∈N. Combining these two cases, we may define *σ* as

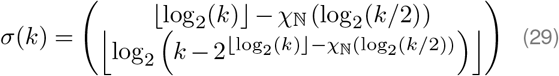

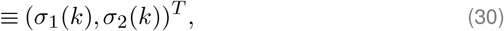

where *χ*_ℕ_ is the indicator function of set ℕ. Hence, Eq. (25) is satisfied. Finally, with *g*(*w*) = 2^*γ*(*w*)^, we define *f* as

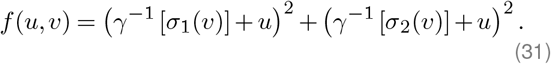

Hence,

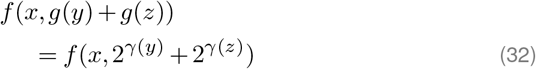

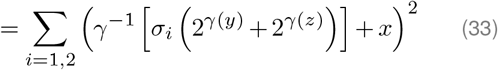

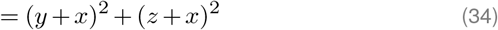

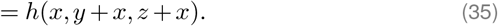

Reverting the change of variables, we have found *f* and *g* such that Eq. (23) holds for all rationals.

We now aim to find “reasonable” examples, in the sense of having some level of literature background and application, of birth and death rates that not only contrast with Champagnat’s framework, but also suggest a potentially more general approach to unifying evolutionary dynamics. Based on these examples, or class of examples, we propose an expansion of the modelling in that paper.

### Evolutionary games on graphs

In evolutionary dynamics, games dependent on cooperativity in a graph structure are often accepted as realistic approximations to describing the dynamics of populations. Thus we now aim to frame evolutionary graph theory in Champagnat’s framework and understand its limitations through examples motivated by other works (Ohtsuki et al., 2006; Allen et al., 2013, 2017).

#### Asymmetric interactions

As discussed below, different degrees and weights of vertices representing individuals might suggest some overall asymmetric growth rates, regarding traits and interactions between specific individuals. For example, the graphs in Figure 2 are a simple first case where Champagnat’s approach could fail, due to the asymmetric nature of the incomplete Graph 2. Considering the focal individual with trait *x*_1_, its dynamics would only be influenced by the individual with trait *x*_2_, as suggested in Graph 2, leading to an interaction kernel equation of the form

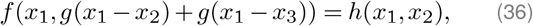

which is not possible due to the symmetry w.r.t. *x*_2_ and *x*_3_ imposed by the left-hand side. Considering the degree of each vertex as an extra trait, however, could solve the mathematical conundrum in Eq. (36), guaranteeing non-focal trait symmetry in some sense, which we discuss next for some examples. Notice also that, even if a specific permutation of the unordered tuple (*x*_2_, *x*_3_) was picked, information on individuals’ labels is required to handle asymmetry, which we discuss in the following.

**Figure 2.**
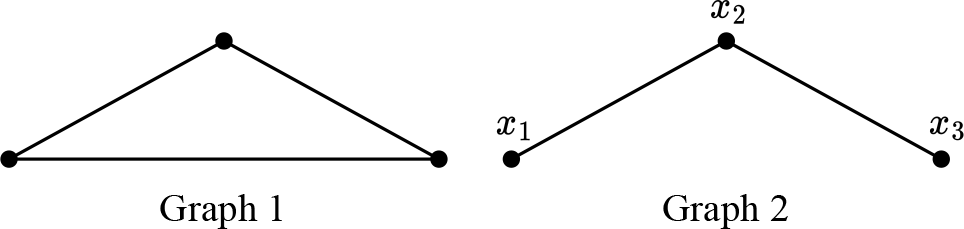
Two evolutionary graphs. In a population in which interaction is defined by incomplete graphs, asymmetry can be overturned by a redefinition of the phenotypic landscape.

#### Incomplete graphs

Consider any graph *G* of order *n* with vertices and edges representing different individuals and their potential interactions, respectively. We assume that the graph is fixed for the duration of the evolutionary dynamics and that the edges are equally weighted. In the one-shot continuous Prisoner’s Dilemma (Poundstone, 1993), a player with cooperativity *x* pays a cost *C*(*x*) to produce a benefit *B*(*x*) for the other player. Here, we consider the game with more players. Therefore, the payoff of a player *i* with cooperativity *x*_*i*_ is

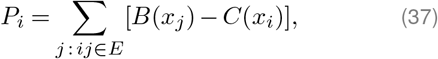

where *E* is the edge set of the graph *G* and *ij* is the edge connecting players *i* and *j*. If *G* is complete, the sum in Eq. (37) is over all *j*. The fitness of player *i, f*_*i*_, is given by a constant term, denoting the baseline fitness, plus the payoff that arises from the game, that is,

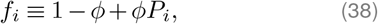

where *ϕ* measures the intensity of selection. For simplicity, and without loss of generality, we consider strong selection, *ϕ* = 1, meaning that the payoff is large compared to the baseline fitness. Hence, *f*_*i*_ = *P*_*i*_. With-out delving into much detail on the biological meaning of fitness and its connection to event rates (Wassersug and Wassersug, 1986; Smith et al., 1989; Bertram and Masel, 2019), we assume that, independently of the chosen update rule (see note below), both birth and death rates of a focal individual *i* depend, in some way, on its fitness. Therefore, taking the cooperativity as a trait, we expect the following dependence

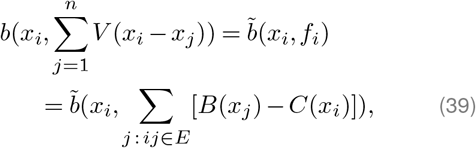

where 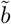 is related to *b* and *V* in a specific way. Similarly for the death rate *d*. The question is now whether *f*_*i*_ is symmetric w.r.t. the other cooperativities *x*_*j*_, *j≠ i*. In general, this is not necessarily true, as seen in the example of Graph 2 (Figure 2), by noting that, for example, *f*_1_ = *B*(*x*_2_) −*C*(*x*_1_)≠ *B*(*x*_3_) −*C*(*x*_1_), ∀*x*_2_, *x*_3_. However, a phenotype redefinition based on the graph’s connectivity could hint at a possible mathematical solution. Notice that, since we have access to the edge set *E*, rewriting Eq. (37) as

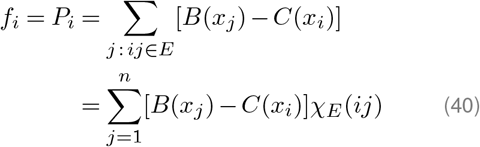

suggests that some extra trait regarding the edge set is needed to guarantee symmetry. Due to Theorem 1, Eq. (39) is always possible as long as Eq. (40) is symmetric w.r.t. non-focal traits. However, the edge set *E* must be somehow obtained from the phenotypes, in order to be consistent with the dependence in Eq. (1)- Eq. (2). In other words, we want to define the set {*j* : *ij*∈*E*} in terms of a certain trait of the phenotype. We now question whether there is a way of relabelling the graph such that, for any vertex *i*, with corresponding new label *p*_*i*_ ∈ ℕ, there exists a set *L* for which

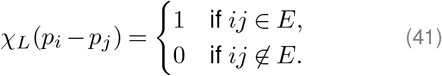

The most straightforward thing to do is take the labels *p*_1_, …, *p*_*n*_ to be so far apart that no difference *p*_*i*_−*p*_*j*_ is repeated, which condition is reminiscent of a *Sidon sequence* (Erdos and Turán, 1941; O’Bryant, 2004). Indeed, we can take *p*_*i*_ = 2*i*. If 2^*a*^−2^*b*^ = 2^*c*^−2^*d*^, then we also have 2^*a*^ + 2^*b*^ = 2^*c*^ + 2^*d*^, and by the uniqueness of binary representations, we must have *a* = *c* and *b* = *d*.Now, let

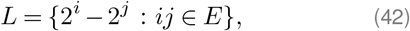

where *E* is the edge set of the graph. Then Eq. (41) holds. Hence, considering *p*_*i*_ = 2^*i*^ as a second trait in the phenotype, we can rewrite Eq. (40) as

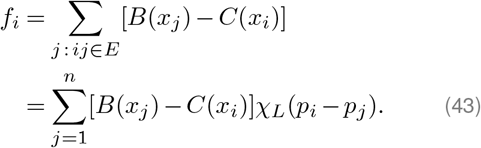

Therefore, it becomes clear that fitness *f*_*i*_, written as a function of two traits (cooperativity and vertex degree), is symmetric w.r.t. similar non-focal traits and thus Champagnat’s framework can be extended to incomplete graphs. Going back to the example Eq. (36), *h* can now be redefined as

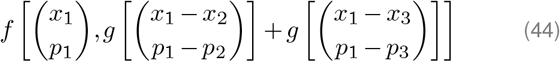

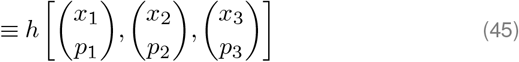

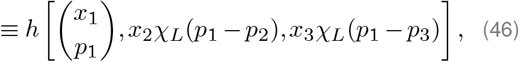

which works for any other focal individual. More generally, we may consider weighted graphs.

#### Weighted graphs

In Allen et al. (2017), the population structure is represented by a weighted, undirected graph *G* with edge weights *w*_*ij*_. The *weighted degree* of a vertex *i* is defined as *w*_*i*_ =∑*j*_∈_*G w*_*ij*_. These weights determine the frequency of game interaction and the probability of replacement between vertices *i* and *j*, considering a death-birth update rule. Here, the notion of cooperation in a game is simplified to two types of individuals, characterised by

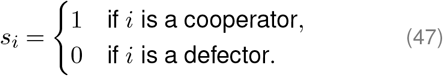

Again, we can think of *s*_*i*_ as a trait. Considering the donation game, the payoff matrix is defined as

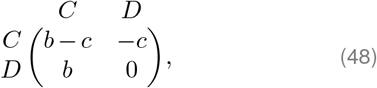

where *C* and *D* represent cooperators and defectors, respectively, and *b, c >* 0 are the benefit and cost values. Then, the edge-weighted average payoff of vertex *i* is given by

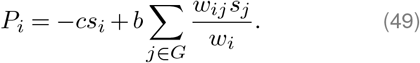

The fitness of individual *i*, which is interpreted as birth rate, is then given by *f*_*i*_ = 1 + *δP*_*i*_, where *δ >* 0 quantifies the strength of selection. Since the weights *w*_*ij*_ are fixed (or interpreted as extra traits), we have that, as long as at least two weights are different, Eq. (49) is not symmetric w.r.t non-focal types *s*_*j*_. Hence, evolutionary dynamics on weighted graphs are also not covered by Champagnat’s framework, unless we consider, again, the connectivity between individuals as traits, which relaxes the definition of asymmetry.

We have then concluded that, from an evolutionary graph theory perspective, the framework in Champagnat et al. (2006) is restricted to symmetric games on well-mixed populations, unless some assumption is made on the phenotype definition to include the edge set asymmetries of the interaction graph.

#### A note on update rules

Evolutionary dynamics depend on update rules upon which these games are settled. In Ohtsuki et al. (2006), for example, a few update rules are considered: death-birth, birth-death and “imitation” updating. However, it may be quite a challenge to adapt them in Champagnat’s framework. The key difference is that Champagnat uses the values of a sequence of random variables uniformly distributed in [0, 1] to select the type of birth or death events, based on the event rates, a methodology not that dissimilar to Gillespie’s algorithm (Gillespie, 1976, 1977). Therefore, they consider a birth-mutation-death process that follows no particular order and where there is no explicit connection between two consecutive events, which may, in some sense, be more realistic. Setting extremely high or low event rates in an artificial manner and introducing specific traits for positioning and number of neighbours might approximate common update rules in evolutionary graph dynamics. Nonetheless, our symmetry argument remains independent of this adaptation.

#### Solving asymmetry

In the previous examples, we have shown that the inclusion of a structure-dependent trait in the phenotypic landscape of the population allows for an adjustment of asymmetric frameworks to be accounted for by Champagnat’s symmetric framework. While seemingly a very artificial argument, incomplete graphs revealed the necessity for the existence of a well-defined trait based on the graph portrait of interaction, in order to guarantee symmetry. The *a priori* knowledge of the interaction structure is key in order to apply the main framework. We call this trait addition the *Sidon-extension* of non-focal phenotypes to ℝ^*ℓ*+1^.

The previous examples Eq. (37) and Eq. (49) are but a small class of very specific *linear* asymmetry. We will see, however, that any asymmetric functions w.r.t non-focal traits can still fit into Champagnat’s framework under a suitable transformation.

##### Example 5

(Symmetrizations). Consider the following asymmetric and non-linear function *h*(*y, z*). In general, such a function could not be represented by an interaction kernel as in Champagnat et al. (2006). However, a symmetrization of *h* based on the relative positioning of individuals (or phenotypes) would extend the previous arguments to any asymmetric function. Pairwise adding extra positional traits *p*_*z*_ *> p*_*y*_, we may write a symmetric version of *h* as follows

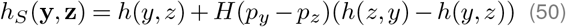

where **y** = (*y, p*_*y*_)^*T*^, **z** = (*z, p*_*z*_)^*T*^, and *H* is the Heaviside step function. Indeed *h*_*S*_ is now symmetric w.r.t. **y** and **z**, and Theorem 1 applies. A relatively straight-forward generalisation to any *n* and *ℓ* is then possible. Alternative and general symmetrization techniques (e.g., Steiner symmetrization) could further refine the definition of *h*_*S*_ and consequent applicability of Champagnat’s framework to general functions (Brock, 1995; Brock and Solynin, 2000). For the purpose of this work, however, we confine our approach to a simple application of boolean functions to construct our generalisation.

An important note is that, while the primary phenotypic space in ℝ^*ℓ*^ fails in the case of incomplete or weighted graphs, its Sidon-extension guarantees the symmetry imposed by the interaction kernels. This is only possible because we are considering vector-wise (or, individual-based) symmetry w.r.t. non-focal traits, and the inclusion of a multiplicative boolean factor in the definition of a general *h*. Without this assumption, Champagnat’s framework generally fails in asymmetric scenarios. We may then write a generalised version of Theorem 1.

##### Theorem 3 (Symmetric asymmetry)

*Assume the Axiom of Choice, and let*

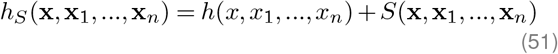

*for some symmetrizing function S, where* **x**_*i*_ ∈ℝ^*ℓ*+1^ *is a Sidon-extension of vector x*_*i*_ *with unique parameter p*_*i*_,∀_*i*_*Then, for all x, x*_1_,.., *x*_*n*_∈R^*ℓ*^ *and n*≥2, *there exist functions f, g and S such that*

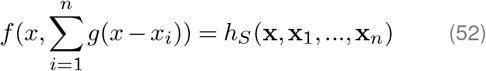

*holds for any function h*.

*Proof*. Let *h* : ℝ^*n*+1^×ℝ^*ℓ*^→ℝbe any function. Take {*p*_*i*_} to be a Sidon sequence (labelling traits) and consider the list of all distinct pairwise combinations of *p*_*i*_ -values. There will be 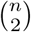 such pairs. For each pair (*p*_*i*_, *p*_*j*_), we define a switch term that accounts for the difference between the original function and the function with *x*_*i*_ and *x*_*j*_ switched. If the original function is *h*(*x, x*_1_, …, *x*_*n*_), the switch term for (*x*_*i*_, *x*_*j*_) maps *x*_*i*_→*x*_*j*_ and *x*_*j*_→*x*_*i*_. By using Heaviside step functions *H*(*p*_*i*_−*p*_*j*_) for each pair, we determine whether or not to apply the switch term. Finally, summing up all these terms weighted by their respective Heaviside functions we guarantee the existence of a function *S* that symmetrizes *h*_*S*_, for any *h*. The existence of *f* and *g* is then a direct consequence of Theorem 1. ◼

## Discussion

In this work, we have explored the realms of logic and set theory to extrapolate an analysis of the symmetry of the interaction kernel definition within a well-established model of evolutionary dynamics (Champagnat et al., 2006).

By invoking the Axiom of Choice, and utilising a mathematical discourse grounded in Zorn’s Lemma, we secured the existence of a Hamel basis. The bijection of this basis to the real numbers enabled us to uniquely determine the sum terms in the dependence of the interaction kernel. The inherently non-constructive nature of this approach further propelled our efforts to examine the symmetry argument across countable sets. This analysis ultimately culminated in two versions of our central theorems pertaining to the overall symmetry of interaction kernels in relation to non-focal traits within an individual’s phenotype (Theorems 1 and 2).

This revelation led us to infer that the framework outlined in Champagnat et al. (2006), although limited to birth and death rate functions that maintain symmetry w.r.t. non-focal traits, can be adapted to accommodate any form of symmetry. Essentially, we illustrated that the event rates could indeed adopt any symmetric functions. Nevertheless, the feasibility of constructing such functions is heavily contingent on the trait values – failing in the real case but possible, in general, within any countable subset of ℝ^*ℓ*^. In the latter scenario, we presented a practical exemplification in the case of nonlinear symmetry.

Ultimately, we ventured into an examination of the symmetry of evolutionary games on graphs. We analysed incomplete and weighted population dynamics graphs in cooperative games, as well as the case of general individual-based asymmetric interaction. We concluded that symmetry can be ensured, provided that individual labelling is incorporated as a component of the phenotype of different individuals (Theorem 3).

Our exploration into the subtleties of symmetry within interaction kernels and the application of set theory notably enhances the modelling and understanding of evolutionary dynamics. By uncovering the roles and limitations of symmetry in shaping interactions between individuals and phenotypic variations, we open a new pathway for a deeper and more nuanced understanding of biological systems. The set theory framework further contributes to the mathematical foundations of these models, allowing for more sophisticated and detailed interpretations. This research serves as a crucial stepping stone in bringing about more refined and predictive models of evolutionary processes, which will be instrumental in furthering our understanding of adaptive evolution and cooperative behaviours within complex biological networks.

## ACKNOWLEDGEMENTS

We thank Prof. Nadia Sidorova and Dr. Rubén Pérez-Carrasco for their helpful discussions and comments that greatly improved the manuscript. This research was made possible by a UCL Studentship Award from the Department of Mathematics and the Leverhulme Trust Research Project Grant RPG-2022-028.

