## Supplementary Information for "Unifying evolutionary dynamics: a set theory exploration of symmetry and interaction"

### A note on Zorn's Lemma

In this section, we sketch a proof that $\mathbb{R}$ has a basis, under the Axiom of Choice and following Zorn's Lemma, and discuss the existence of the critical bijection $g$ between $\mathbb{R}$ and its Hamel basis. See Halpern (1966); Bleicher (1967) for further reading.

First, we recall some definitions from set theory (Harzheim, 2005)

- A *partially ordered* set is a set $X$ together with an ordering $\leq$ of the elements of $X$ that is transitive (if $x \leq y$ and $y \leq z$ then $x \leq z$) and antisymmetric (if $x \leq y$ and $y \leq x$ then $x = y$);
- A *totally ordered* subset of $X$ is a set $Y \subseteq X$ such that, $\forall y, z \in Y$, either $y \leq z$ or $z \leq y$;
- A *chain* $Y$ in a partially ordered set $X$ is a totally ordered subset of $X$;
- An *upper bound* for a subset $Y$ of a partially ordered set $X$ is an element $u$ such that $y \leq u$ for every $y \in Y$;
- Finally, a *maximal element* in a partially ordered set $X$ is an element $x_0$ such that the only element $x \in X$ satisfying $x_0 \leq x$ is $x_0$ itself.

The main result follows.

**Theorem 4** ($\mathbb{R}$ has a basis)**.** *Under the Axiom of Choice, there exists a basis of* $\mathbb{R}$.

*Proof.* (Sketch) Recall that $\mathcal{B}$ is defined as the collection of real numbers such that every real number is a rational linear combination of numbers $\beta_i \in \mathcal{B}$ in precisely one way, that is, every real number $\alpha$ has a unique representation of the form

$$\alpha = \sum_{i=1}^{m} q_i \beta_i, \tag{S1}$$

where $q_i$ is rational and $m$ depends on $\alpha$. It suffices to show that $\mathcal{B}$ exists.

With $\mathcal{B}$ as a candidate $\mathbb{Q}$-basis of $\mathbb{R}$, the objects we are interested in are subsets of $\mathbb{R}$ that are linearly independent over $\mathbb{Q}$. First, we note that a maximal linearly independent subset of $\mathbb{R}$, where by "maximal" we mean that the set is not contained in any larger linearly independent set, spans the whole of $\mathbb{R}$. If it did not, we could just pick an element that did not belong to its linear span and add it to the linearly independent set, redefining maximality. Although seemingly an interminable process, Zorn's lemma tells us that it ends. Thus, we are looking at the set of all linearly independent subsets of $\mathbb{R}$, with the partial order $\subset$.

In order to apply Zorn's Lemma, we then need to check that every chain has an upper bound. Imagine that we have a collection $Y$ of linearly independent subsets of $\mathbb{R}$ and that, for any two of those subsets, one is contained in the other. By definition, the upper bound of such a chain is a set that contains all the sets in $Y$, so it has to contain their union. If we take the upper bound as the union, it should be linearly independent. Indeed, if $\alpha_1, \ldots, \alpha_n$ belong to the union, then each $\alpha_i$ belongs to some linearly independent set $\beta_i \in Y$. Since $Y$ is a chain, one of these sets $\beta_i$ contains all the others. If that is $\beta_j$, then the linear independence of $\beta_j$ implies that no non-trivial linear combination of $\alpha_1, \ldots, \alpha_n$ can be zero, which proves that the union of the sets in $Y$ is linearly independent. Therefore, by Zorn's Lemma, there is a maximal linearly independent set. Such a set forms a basis for $\mathbb{R}$. ■

Now, in order to define $g$ in Eq. (3) as the bijection between $\mathbb{R}$ and its Hamel basis, it is important to show that such a basis has the cardinality of the continuum. Let the cardinality of some Hamel basis be $\kappa$. Since the cardinality of the set of coefficients is $\aleph_0$ (cardinality of $\mathbb{Q}$), we have $\aleph_0\kappa$ elements using each basis element of the form $q_1\beta_1$, $\aleph_0^2\kappa^2$ elements using two basis elements $q_1\beta_1 + q_2\beta_2$, and so on. Hence, the span of the basis has cardinality given by

$$\aleph_0\kappa + \aleph_0^2\kappa^2 + \aleph_0^3\kappa^3 + \cdots = \kappa, \tag{S2}$$

where we used the fact that $\aleph_0$ is the smallest infinite cardinal (the product of two infinite cardinals is equal to whichever is bigger, thus $\aleph_0\kappa = \kappa$). Since the span is $\mathbb{R}$, we have $\kappa = 2^{\aleph_0}$ and the bijection exists.

Egbert Harzheim. *Ordered sets*, volume 7. Springer Science & Business Media, 2005.
